# Asymmetric adaptation reveals functional lateralization for graded versus discrete stimuli

**DOI:** 10.1101/411033

**Authors:** Melanie Desrochers, Marianne Lang, Michael Hendricks

**Affiliations:** Department of Biology, McGill University, Montreal, Quebec H3A 1B1, Canada; École normale supérieure de Lyon

## Abstract

Animal navigation strategies depend on the nature of the environmental cues used. In the nematode *Caenorhabditis elegans*, navigation has been studied in the context of gradients of attractive or repellent stimuli as well is in response to acute aversive stimuli. We wanted to better understand how sensory responses to the same stimulus vary between graded and acute stimuli, and how this variation relates to behavioral responses. *C. elegans* has two salt-sensing neurons, ASEL and ASER, that show opposite responses to stepped changes in stimulus levels, however only ASER has been shown to play a prominent role in salt chemotaxis. We used pre-exposure to natural stimuli to manipulate the responsiveness of these neurons and tested their separate contributions to behavior. Our results suggest ASEL is specialized for responses to acute stimulus changes. We also found that ASER remains responsive to graded stimuli under conditions where it is unresponsive to large steps.

## Introduction

Organisms ranging from bacteria to multicellular animals use conserved navigation strategies to locate resources in their environment despite enormous differences in the specific biological implementation of these strategies. Navigation strategies are specific to the spatial or temporal features of the substances that constitute this information. One of the most common types of navigation relies on sensing and tracking gradients of diffusing substances. Depending on the shape and size of these gradients, the relevant sensors—whether they are receptors of a bacterium or axonal growth cone or the olfactory apparatus of a mammal—must continually undergo cycles of sensitization and desensitization to remain responsive to small changes in the stimulus. This is potentially challenging, because the concentration of an attractant my change over orders of magnitude. At the same time, animals must remain sensitive to abrupt changes in stimuli that may require acute behavioral responses.

In *Caenorhabditis elegans,* salt chemotaxis has been used to dissect navigation strategies used at the genetic, cellular, circuit, and behavioral levels. The primary salt sensing neurons in *C. elegans,* ASEL and ASER, exhibit opposite responses to stepped changes in salt concentration: ASEL is activated by cations and responds to up-steps in salt concentration, while ASER detects anions and responds to down-steps (Pierce-Shimomura et al., 2001; Suzuki et al., 2008). Under most conditions, gradient chemotaxis in the nematode *Caenorhabditis elegans* can be described as a biased random walk, a strategy identical to that used by single-celled organisms like bacteria, multicellular growth processes observed in plant roots, and even subcellular structures like axonal growth cones (Berg and Brown, 1972; Pierce-Shimomura et al., 1999). While *C. elegans* does respond to salt up-steps and down-steps (Miller et al., 2005), most behavioral analysis is done in the context of gradient navigation. However, the physiological properties of ASEL and ASER are less well-explored in the context of graded stimuli. ASER shows complex calcium responses to gradients that depend both on the direction of salt change and whether salt levels are above or below a memorized cultivation concentration (Luo et al., 2014). While ASEL contributes to forward locomotion during exposure to attractive salt concentrations (Suzuki et al., 2008), ASEL has been found to be largely dispensable for both positive and negative salt gradient chemotaxis (Kunitomo et al., 2013; Luo et al., 2014), though recently it was shown that optogenetic stimulation of ASEL can drive fictive positive chemotaxis under specific conditions (Wang et al., 2017).

We were interested in attempting to reconcile calcium imaging experiments, usually performed using stepped stimuli, with behavioral studies that examine behavior in stimulus gradients. The increasing use of microfluidics for both calcium imaging and behavioral analysis provides an opportunity to do so (Albrecht and Bargmann, 2011; Chronis et al., 2007), though producing reproducible temporal stimulus gradients in microfluidic devices is challenging. We previously showed that gradients could be generated by coupling a mixing chamber with magnetic stir plate (Luo et al., 2014). However, this system was cumbersome and difficult to control.

Here, we describe a dedicated device based on the same principle that can deliver reproducible temporal gradients into microfluidic devices for both calcium imaging and behavioral experiments. We show that ASEL and ASER can be separately and selectively desensitized or sensitized by pre-exposure to high or low salt concentration and use this manipulation to test the contribution of each to behavioral responses to either stepped or graded stimuli. The use of real stimuli to manipulate sensory neuron sensitivity has the advantage of not relying on artificial manipulations of neural circuit function, thus the responses and behaviors we observe all fall within the normal physiological and functional repertoire of the animal. We find that ASEL sensitization enhances behavioral responses to stepped stimuli. ASER-sensitized animals ignore large steps despite showing robust calcium responses to the stimulus. In response to graded stimuli, ASER desensitization specifically impairs response to salt increases, suggesting that even when ASEL is sensitized it is not sufficient to drive chemotaxis in our assay. Finally, we observe that under conditions in which ASER is desensitized and unresponsive to salt down-steps, it remains sensitive to negative salt gradients, exhibiting sharp, sporadic calcium events, and the probability of these events is regulated by integrating graded salt decreases over time.

## Materials and methods

*Caenorhabditis elegans* were raised on nematode growth medium, 0.25% Tryptone (w/v), 1.5% Agar (w/v), 1 mM CaCl2, 1 mM MgSO4, 25 mM KPO4 (pH 6.0), 50 mM NaCl, and fed with *E. coli* OP50 (Brenner, 1974; Stiernagle, 2006). Young adult hermaphrodites were used in all experiments. Strain ZC1600 [*yxEx783*], previously described, contains an extrachromosomal array driving expression of the genetically-encoded calcium indicator GCaMP3 under the *flp-6* promoter (Luo et al., 2014; Tian et al., 2009).

### Salt solutions

Salt solutions were made in a buffer identical to nematode growth medium (NGM) except without agar, tryptone, or cholesterol, 1 mM CaCl2, 1 mM MgSO4, 25 mM KPO4 (pH 6.0).

### Temporal gradient production

We designed an instrument to produce temporal gradients based on rapidly mixing small volumes of two solutions off-chip prior to flowing into the stimulus channels or behavioral arena. Schematic and code used to control the device are found in Figure 3 and Supplementary Figure S1. Briefly, we made two solutions with different concentrations of salt and fluorescein (to monitor the gradient) corresponding to the highest and lowest concentration for a particular stimulus sequence. Solenoid pinch valves (Neptune Research, 225P011-21) controlled the flow of these solutions into a small mixing chamber made from 0.25 inch ID tubing containing either a 7 mm magnetic stir bar or a small spherical magnet (Fisher Scientific). A larger stir bar was affixed to a 12V DC motor and the mixing chamber was positioned close to it, clamped in place with an alligator clip. Driving the motor to spin the large bar causes rapid mixing of solutions in the chamber. The DC motor and valves were controlled by an Arduino UNO R3 microprocessor (see Supplemental Figure S1). Gradient steepness and shape is a function of flow rate, mixing chamber volume, and pinch valve duty cycles. The lag between switching solutions and affecting the in-chip gradient can be reduced by positioning the mixing chamber as close as possible to the microfluidic device. Because the fluorescein salt is itself an ASE stimulus, it was used over a concentration range 4to 6 orders of magnitude smaller than the NaCl range and in the opposite polarity, such that the highest fluorescein concentration (0.08 μM for calcium imaging, 1 μM for behavior) corresponded to the lowest salt concentration and vice versa.

### Microfluidic device fabrication

Photoresist (SU8) masters for microfluidic devices were fabricated by soft lithography (San-Miguel and Lu, 2013). Polydimethylsiloxane (PDMS) (Dow Corning Sylgard 184, Ellsworth Adhesives #4019862) was mixed at 10:1, degassed, poured over masters, degassed again, and cured at 60°C for at least 3 hours. Devices were replica mastered in a two-part epoxy resin (Smooth Cast 310, Sculpture Supply Canada #796220). Inlet holes were bored with a Milltex 1 mm biopsy punch (Fisher). Chips were cleaned and bonded to glass coverslips using air plasma generated by a handheld corona treater (Haubert et al., 2006) (Electro-Technic Products, Chicago, IL). Coupling to fluid reservoirs was done by directly inserting PTFE microbore tubing (Cole-Parmer #EW-06417-21) into inlet holes.

### Calcium imaging

Fluorescence time lapse imaging was performed on an Olympus IX83 inverted microscope using a 40x silicone immersion lens. Images were captured with an OrcaFlash4.0 sCMOS camera (Hamamatsu). A piezo-controlled z-axis stage (Prior Scientific) was used for alternately imaging ASEL and ASER by rapidly switching between focal planes. Because of the long time course of the experiments, animals were imaged at low illumination levels using 300 ms exposures and capture each neuron at 1 Hz. Animals restrained in a microfluidic channel (Chronis et al., 2007) were exposed to the stimulus streams described in each experiment. For stepped stimuli, fluid flow was controlled with a ValveBank (AutoMate Scientific) and driven by a vacuum trap on the outflow channel. Graded stimuli were produced by the off-chip mixing device described in Figure 3, Supplemental Figure S1, and below. Fluorescence intensity was measured from neuronal soma using Fiji (Schindelin et al., 2012). For Ca^2+^ traces and statistical analysis, fluorescence was normalized to the first 3 seconds of recording, for heat maps intensities were normalized to the minimum value for each animal.

### Behaviour tracking

Animals were loaded into a microfluidic device based on the "pulse chip" (Albrecht and Bargmann, 2011). The chip consists of an array of posts sized and spaced to allow *C. elegans* to crawl through a fluid substrate under constant, controllable flow. For each assay 20-30 animals were loaded into the microfluidic arena after pre-exposure and presented with stimulus sequences described for each experiment while being recorded with a USB camera (Logitech 210 or Dino-Lite). Fluid flow was produced by gravity. Position over time for each animal was measured with the TrackMate ImageJ plugin (Tinevez et al., 2017). Tracks shorter than 60 seconds were excluded. Frames where animals explored the chip boundary were excluded. From these coordinates, a path angle measure was extracted for each frame by calculating the angle between vectors projecting from the current animal position to future and past positions with a 5 second lag.

### Data analysis

JMP Pro 13 (SAS) was used for statistical tests and graphs.

## Results and Discussion

Sensory systems adapt based on a recent experience. Sensitization is expected to occur when stimulus levels are low, and desensitization is expected with high stimulus levels. Because ASEL and ASER respond to salt steps of opposite valence, high or low salt concentrations should have opposing effects on adaptation in these two neurons. We explored the effects of prior exposure to higher or lower salt on ASER and ASEL calcium responses by removing animals from food and washing them in a buffer containing either 0 mM or 50 mM NaCl for 15 or 60 minutes (Figure 1A).

**Figure 1.**
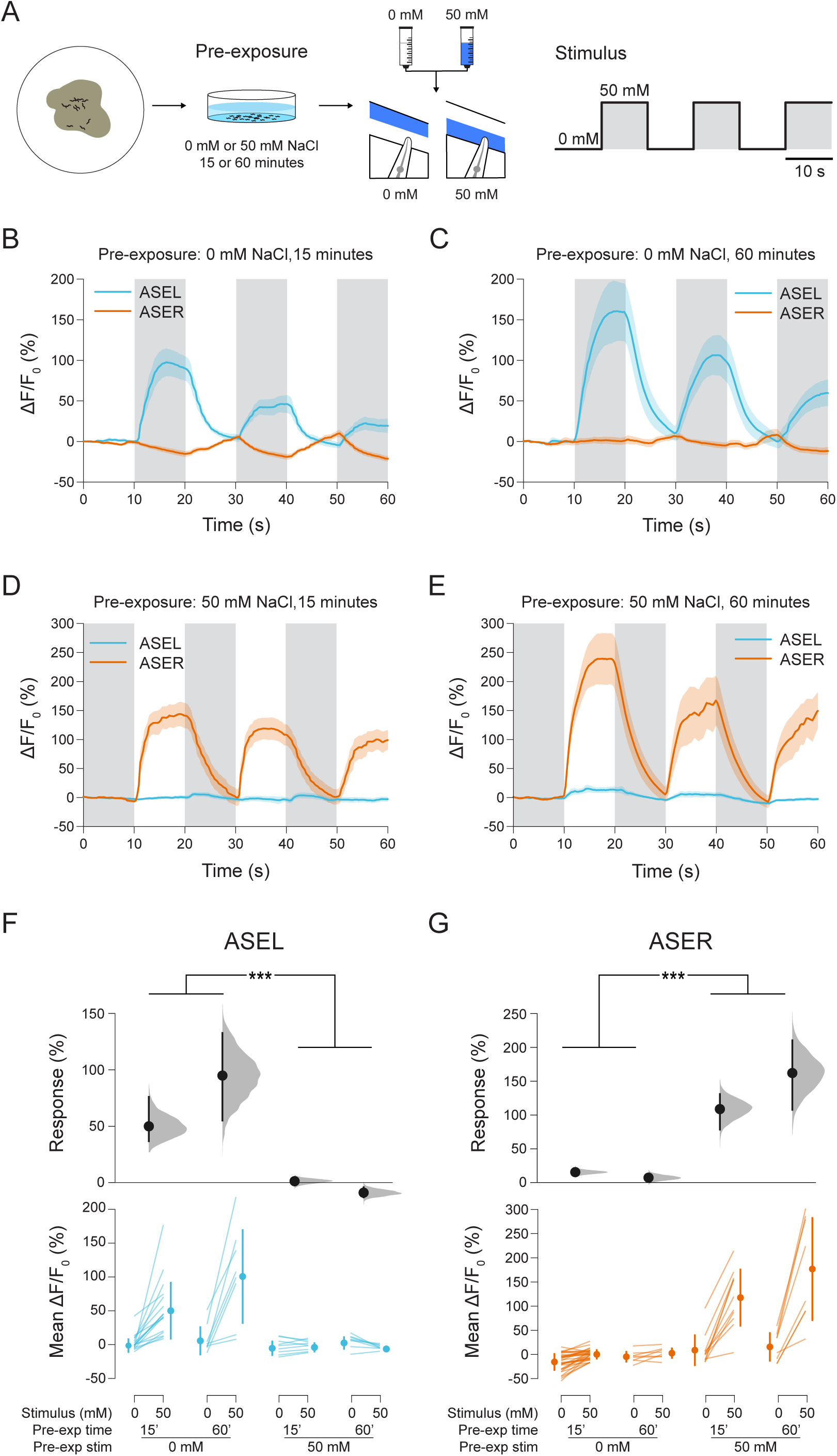
Modulating ASEL and ASER responsiveness with stimulus pre-exposure. (A) Animals were raised under standard conditions. Prior to calcium imaging, animals were washed in a solution containing either o mM or 50 mM NaCl for 15 or 60 minutes prior to imaging. (B) Pre-exposure too mM NaCl attenuates ASER responses, while ASEL shows robust ON responses. (C) Longer pre-exposure increases ASEL response magnitude. (D) Pre-exposure to 50 mM NaCl attenuates ASEL responses while ASER shows strong OFF response. (E) One hour of pre-exposure increases ASER responses. (F) Comparison of mean fluorescence intensity in the 5 second time windows shown in yellow in (A) by pre-exposure condition and stimulus in ASEL. (G) Comparison of mean fluorescence intensity in the 5 second time windows shown in yellow in (A) by pre-exposure condition and stimulus in ASER. Sample size for each condition left to right ASEL/ASER n = 17/33, 8/8, 9/11, 8/8. A two-way ANOVA to test the effects of pre-exposure condition and time on response magnitude was significant for ASEL (F_3,3s_ = 13-3386, *P* < 0.0001, significant main effect of conditionP < 0.0001) andASER (F_3,56_ = 45-7473, *P* < 0.0001, significant main effect of conditionP < 0.0001). Pre-exposure time was not significant. ***P < 0.001, post-hoc Student’s t-test.

Animals expressed the genetically-encoded calcium indicated GCaMP3 in both ASE neurons. After pre-exposure, animals were loaded into a microfluidic device that exposes the nose of restrained animals to alternating fluid streams (Chronis et al., 2007). We recorded GCaMP fluorescence intensity in ASER and ASEL while animals were exposed to 50 mM steps in salt concentration. We observed the well-characterized ON (ASEL) and OFF (ASER) properties of the two neurons in response to stepped stimuli, however we found that their responses were dramatically altered by pre-exposure to high or low salt concentrations (Figure 1B-E). ASEL, which responds to salt upsteps (Suzuki et al., 2008), showed minimal responses to a 5o mM upstep in salt concentration when animals were pre-exposed to high salt (adaptation), but large responses when pre-exposed to a salt-free buffer (sensitization). ASER exhibited the opposite pattern of adaptation/sensitization. Both short (15 minute) and long (60 minute) pre-exposures had similar effects (Figure 1F,G).

We reasoned that this asymmetric effect could be used to test the contribution of each of these neurons on behavioral responses to subsequent stimuli. Animals were exposed to alternating pulses of high and low salt solutions in a microfluidic arena (Albrecht and Bargmann, 2011). We used a simple measure of path geometry to quantify behavioral responses (Pradhan et al., 2018) (Figure 2A). Lower path angles indicate forward crawling, associated with preference for a stimulus; high path angles are associated with frequent reversals, turns, and slower locomotion, behaviors typically associated with avoidance.

**Figure 2.**
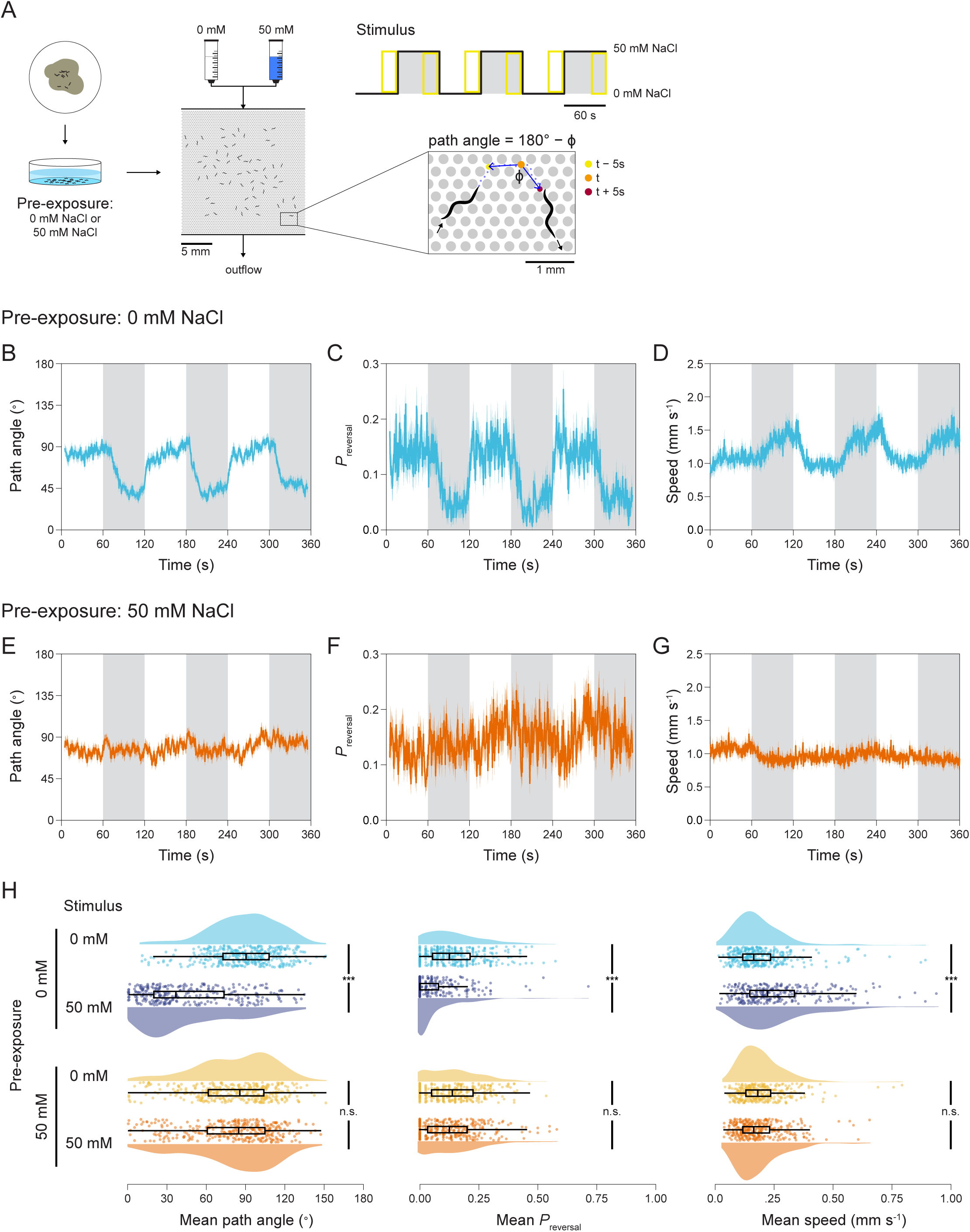
Behavioral responses to pulsed stimuli are affected by pre-exposure. (A) Animals were raised and subjected to pre-exposure conditions as in Figure 1 then loaded into a microfluidic arena where they were exposed to alternating stimulus steps of o mM and 50 mM NaCl in buffer. Behavior was quantified by measuring the speed and path angle of each animal (see Methods). A path angle< 10° was defined as a reversal. (B-D) Animals pre-exposed too mM NaCl showed robust behavioral responses to steps between o mM and 50 mM NaCl in path angle, reversal state probability, and speed. (E-G) Animals pre-exposed to 50 mM NaCl showed no behavioral responses to step stimuli. Traces are mean and shading is standard error. (H) Quantification of traces shown in (B-G) showing distribution (kernel density estimation), quartiles, and points corresponding to each track. Means for each measurement were determined for each track over a 20-second time window corresponding to the yellow boxes in (A). For tracks spanning more than one window, a single mean over multiple windows was calculated. ***P < 0.001 Student’s t-test. Because of tracking errors, collisions, and edge encounters, sample size varies over the course of the experiment. For 50 mM pre-exposure, n = 142-193, for o mM pre-exposure, n = 126-181.

**Figure 3.**
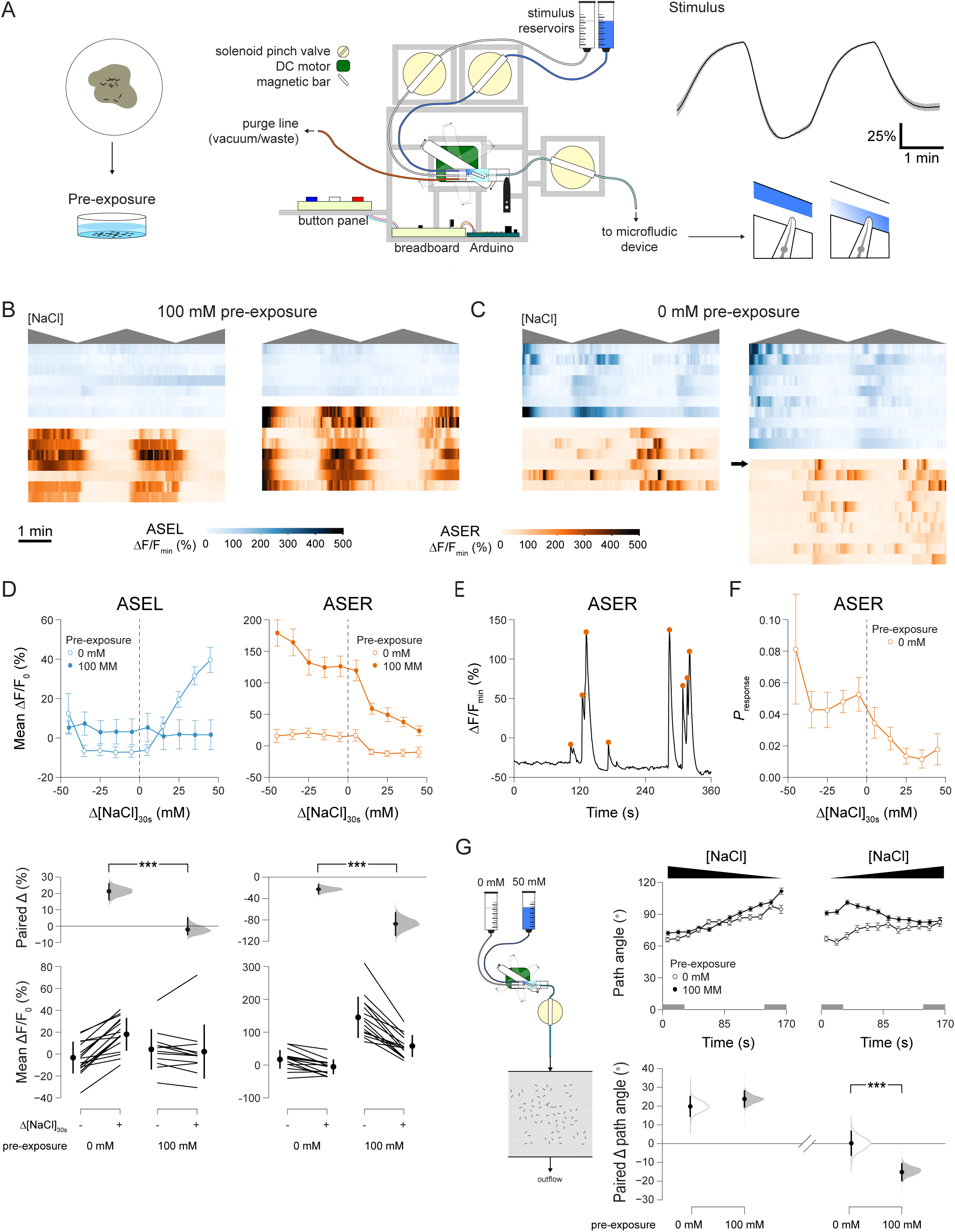
Generation of temporal gradients. (A) Two solenoid valves control the flow of solutions representing the minimum and maximum salt concentration into a mixing chamber. A magnetic stir bar spun by a DC motor drives a small stir bar in the mixing chamber. The mixed solution flows out to a microfludic device. The valves and motor are controlled with a button interface through a programmable microcontroller (see Methods, Supplemental Figure 1). An example of gradient produced by the machine are shown at right, shading is standard error (n = 15). (B) Calcium responses of ASEL (top) and ASER (bottom) to graded stimuli shown as heat maps, gray indicates direction of the salt gradient. (D) Mean GCaMP intensity as a function of the net change in NaCl concentration over the past 30 seconds in ASEL (left) and ASER (right). Responses for each animal in response to positive or negative changes are shown below as estimates of the mean difference (Paired /),., distribution of estimated mean with measured mean and 95% confidence interval) and paired plots (individual measurements, mean and standard deviation). Whether the change in NaCl over the preceding 30 seconds was positive or negative was a significant effect on paired responses for ASEL and ASER. Repeated measures ANOVA, ASEL F1,2_7_ = 35.0174, *P* <0.0001, ASER Fi,2_9_ = 29.5345, *P* < 0.0001. (E) Example of calcium events seen in ASER after pre-exposure too mM NaCL. Orange dots indicate " responses " scored as local maxima occurring after high rates of intensity increase (see methods). (F) Relationship between change in NaCl over the previous 30 seconds and probability of a response, as defined in (E). Gradient direction was a significant effect on response probability. ANOVA F1,_30_ = 14.1617, *P* = 0.0007. (G) Imaging behavior in response to temporal gradients. Changes in mean path angle over time periods of salt decrease (left) and salt increase (right). Mean paired changes in path angle according to pre-exposure treatment (below). Gray shading on time axis indicates time windows used for comparison. *\*\*\*P* < 0.001 paired t-test.

Animal pre-exposed to NaCl-free buffer showed robust behavioral responses to alternating pulses of O mM and 50 mM salt solutions, exhibiting reduced path angles in the preferred, high salt solution and lower path angles in the low salt solution (Figure 1B). This corresponded to changes in reversal frequency (Figure 1C) and speed (Figure 1D). In contrast, animals pre-exposed to 50 mM salt showed no behavioral responses to salt steps between 0 mM and 50 mM (Figure 2E-G). Thus, selective sensitization of ASEL renders animals behaviorally responsive to large salt steps, while sensitization of ASER and desensitization of ASEL results in animals that ignore these steps (Figure 1H).

We next wanted to compare how selective sensitization affected responses to graded stimuli. We designed a compact instrument that controlled fluid flow into microfluidic devices and included a low-volume, in-line mixing chamber, in which a small magnetic stir bar is driven by a larger magnet attached to a DC motor (Figure 1A). The valves and motor were controlled by an Arduino microcontroller (see Materials and Methods and Supplemental Figure S1). This device allows the production of reproducible temporal gradients flowing through the chip.

We first tested how pre-exposure affected ASEL and ASER responses to graded stimuli. We observed that sensitization / desensitization relationships were overall like those for stepped stimuli. Pre-exposure to 100 mM NaCl produced robust and sustained ASER calcium activity in response to negative gradients and highly attenuated, if any, ASEL responses (Figure 3B). Pre-exposure to 0 mM NaCl led to robust ASEL activity in response to positive gradients. Unlike in the stepped-stimulus assay, however, we also observed ASER responses to negative gradients—salt decreases—after this treatment (Figure 3C).

To quantify neuronal responses relative to the gradient, we plotted mean GCaMP intensity relative to recent (30 seconds) net estimated changes in salt concentration (Figure 3D, upper panels) and measured the paired difference for each animal between responses to negative and positive gradients (Figure 3D, lower panels). These findings were consistent with stepped stimuli. However, the short, sporadic responses observed in ASER after 0 mM pre-exposure are not well-captured by an analysis of fluorescence averaged over time. We decided to quantify these as discrete events by detecting local maxima preceded by high rates of intensity increase (Figure 3E). We then calculated the probability of these events relative to amplitude gradient direction (Figure 3F) and found that these events, like sustained responses, are regulated by recent stimulus history.

We next generated temporal gradients within the behavioral microfluidic device. Here, results were quite different from those using stepped stimuli. Pre-exposure to 0 mM or 100 mM NaCl did not affect the ability to adjust behavior in response to a negative salt gradient, indicated by a steadily increasing path angle (Figure 3G, left). In contrast, pre-exposure did influence the ability to respond to negative salt gradients, with animals pre-exposed to 100 mM showing path angle decreases and those exposed to 0 mM showing no change (Figure 3G, right).

Together, these results suggest that ASEL and ASER are specialized for stepped versus graded salt stimuli, respeictivel, and this may explain the lack of any role for ASEL in most salt chemotaxis assays. When ASEL is sensitized, animals show robust changes in locomotion in response to abrupt switches between high and low salt solutions. When ASER activity predominates and ASEL is attenuated, animals ignore these switches. Gradients are more complex. ASEL is not sufficient on its own to mediate positive gradient chemotaxis, as only animals pre-exposed to 100 mM NaCl (ASER-sensitized) show behavioral responses. However, both ASEL- and ASER-sensitized animals can navigate in response to a negative temporal gradient. We suggest that this is because, despite ASER being unresponsive to stepped stimuli after 0 mM pre-exposure, it remains responsive to graded stimuli through a different mode of activity characterized by sharp, discrete calcium events rather than long sustained calcium waves.

While it is ideal to measure activity and behavior at the same time, this is often not possible, and precisely controlling stimuli is difficult in freely-moving animals. With the exception of temperature, the attention paid to *C. elegans* behavior to stimulus gradients has not been matched by analysis of neural responses to graded stimuli, and it is clear there are important differences in responses to graded versus stepped stimuli (Itskovits et al., 2018; Larsch et al., 2015; Luo et al., 2014). Our approach overcomes this technical barrier to allow easy analysis of neuronal responses to gradients. We also show that ASER shows strikingly different patterns of activity depending on stimulus regimen and prior exposure. It was recently reported that the AWA olfactory sensory neuron also exhibits diverse physiological modes depending on context (Liu et al., 2018). These observations, combined with our use of natural stimuli to manipulate sensory responses, underscore the need to analyze neuronal function across their physiological ranges in response to natural stimuli in order to understand sensorimotor transformations.

## Acknowledgements

We thank Xinyu Liu and Xianke Dong (McGill University) for providing photoresist masters for microfluidic devices. We thank all members of the lab for helpful discussions and comments. Some strains were provided by the Caenorhabditis Genetics Center (CGC), which is funded by NIH Office of Research Infrastructure Programs (P40 OD010440).

## Author Contributions

M.D. and M.H. designed the gradient instruments. All authors designed experiments, M.D. and M.L. conducted experiments. All authors analyzed the data. M.D. and M.H. wrote the manuscript.

---

**Supplemental Figure S1.**
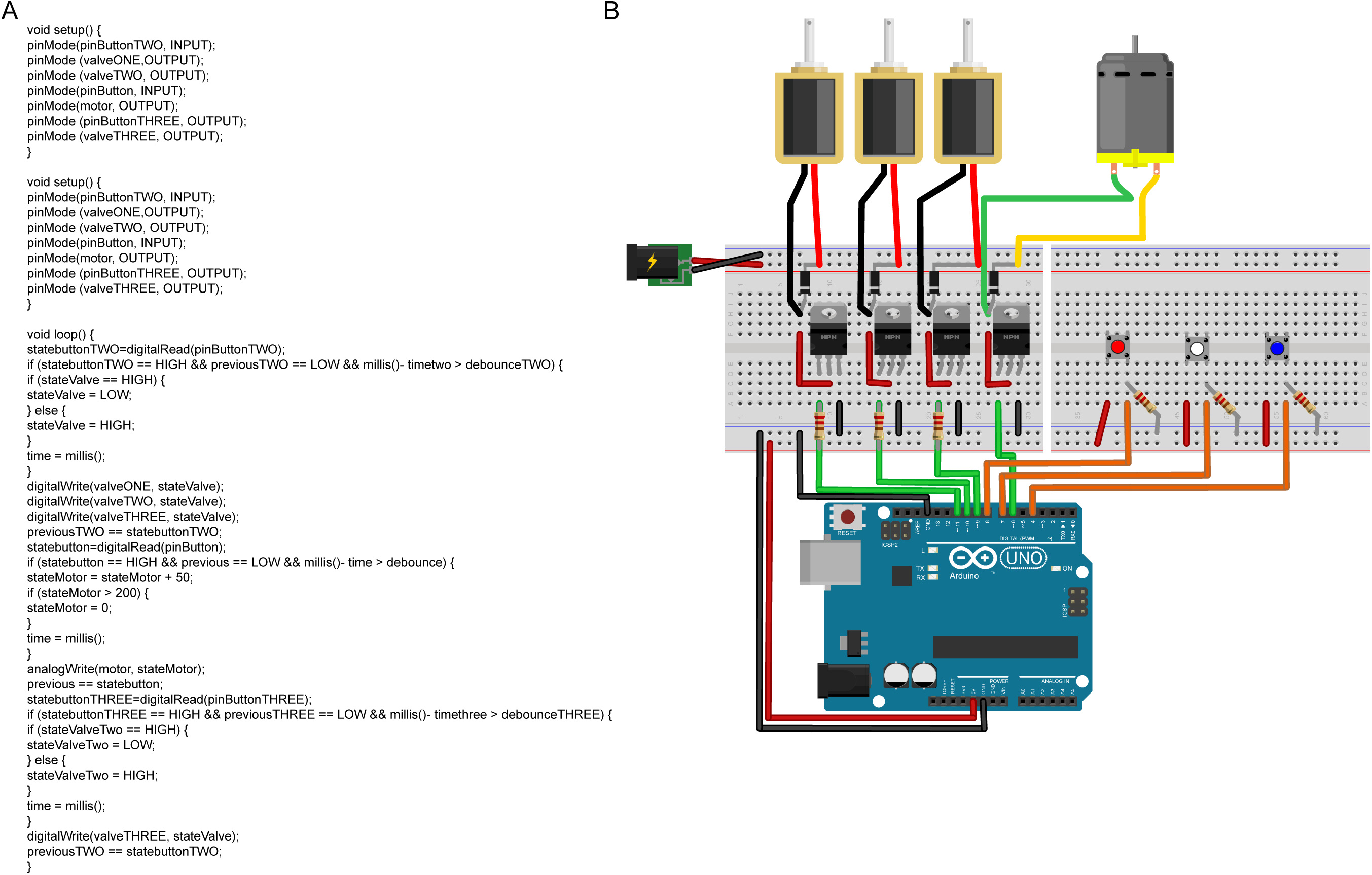

